# FUSION: A fast and uniform processing framework for whole-brain optical scattering tomography images

**DOI:** 10.64898/2026.01.06.698063

**Authors:** Chong Chen, Peilin Gu, Martin Villiger, Néstor Uribe-Patarroyo, Brett Bouma, Jian Ren

## Abstract

Coherent scattering imaging has seen substantial development in clinical ophthalmology and endoscopic microscopy owing to its high throughput, large dynamic range, and capability for label-free imaging. However, limited optical penetration has constrained its application to whole-organ imaging studies. Here, we propose a high-throughput coherent scattering imaging system assisted by tissue clearing, termed *fu*CAST, which enables whole-brain white-matter imaging. In parallel, we develop a high-throughput image reconstruction and uniform stitching platform, FUSION, specifically tailored for *fu*CAST datasets.

## 1. Introduction

Whole-brain, systems-level studies, including connectomic analyses, are critical for understanding neural function and the pathogenesis of neurological disorders. Imaging platforms such as fMOST and serial two-photon tomography integrate optical imaging with physical sectioning to generate cellular-resolution whole-brain datasets and reference atlases that have become de facto gold standards. Nevertheless, unavoidable tissue loss from sectioning, coupled with complex instrumentation and demanding downstream image processing, has constrained their scalability and broad deployment. In recent years, advances in tissue clearing have transformed optical imaging by enabling systematic, organ-wide interrogation of intact tissues, particularly allowing the brain to be studied from a global, systems-level perspective. Concurrently, improved optical penetration has driven the development of optical microscopy techniques for whole-brain imaging, most notably light-sheet microscopy^1^, which in recent years has diversified into numerous implementations enabling imaging of entire mouse brains and even whole mouse bodies. However, a widely accepted and robust method for antibody staining of intact whole mouse brains has yet to be established. Existing approaches typically suffer from either prohibitively long staining times^2^ or require specialized, expensive instrumentation under stringent experimental conditions^3^. Importantly, fluorescence bleaching during prolonged, centimeter-scale three-dimensional imaging poses a fundamental limitation for clearing-based fluorescence microscopy.

Several label-free imaging approaches, such as second-harmonic generation (SHG)^4,5^ and stimulated Raman scattering microscopy (SRS)^6^ have been investigated for imaging cleared brain tissue, yet their low imaging speed and limited throughput fundamentally restrict their applicability to large-scale studies. As a label-free optical imaging method, optical coherent tomography (OCT) within high throughput and high dynamic range has demonstrated its superiority in clinical ophthalmic and vascularendoscopes diagnosis^7,8^. Currently, light scattering in tissues significantly restricts the imaging depth of optical methods, resulting in limited research on OCT imaging of large isolated organs or tissues. Only a few methods based on serial sectioning and invasive microneedling achieved *ex-vivo* deep brain imaging ^9–11^. Losing or damaging the sample means that it is difficult to obtain complete information about the sample using these methods. In our previous research, our lab developed a Clearing Assisted Scattering Tomography (CAST) method and achieved mouse brain hemisphere scattering imaging^12^. The CAST method effectively addresses the imaging depth limitations of OCT while enhancing the signal-to-noise ratio of neural fiber and vascular structures. However, when extended to whole-organ applications, CAST still faces challenges in both imaging throughput and data-processing throughput. The field of view (FOV) of CAST system is limited to just 4 square millimeters. Given the size of the mouse brain (∼11 ×13 × 8 mm^3^), CAST requires multi-dimensional stitching to reconstruct complete whole-brain datasets. Limitations in imaging throughput further compound the complexity of large-scale data stitching. In addition, post-processing speed - including image reconstruction and stitching of large CAST datasets - has become a major bottleneck for large-cohort applications.

Image reconstruction is a complex computational process that requires interpolation, inverse Fourier transform and other operations on a line-by-line basis. Over the past few years, owing to the development of GPU parallel computing technology, several ultra-fast coherent scattering reconstruction algorithms have been developed using hardware acceleration ^13–15^. However, coherent scattering imaging of large *ex-vivo* samples is still in the exploratory stage. To the best of our knowledge, no high-throughput OCT image stitching method optimized for large datasets has been developed yet. In recent years, numerous methods have been developed for large-scale three-dimensional image stitching and striping artifact correction, and have been successfully applied to fluorescence images acquired by light-sheet and confocal microscopy. While there are many fast stitching methods for large datasets of fluorescence images, which have made many attempts in terms of stitching speed and seamless stitching ^16–18^, the existing stitching algorithms cannot be directly applied to stitching CAST images due to the next two reasons. Firstly, the image acquisition mode of CAST is unique. CAST is based on A-Scan as the basic acquisition unit. Data reconstruction and storage is also A-line by A-line. And each A-line data contains both sharp in-focus and low-resolution out-of-focus information. Therefore, the CAST image must accurately extract the in-focus portion of each A-line data prior to stitching. At the same time, the extra acquisitions of A-line data further increase the size of the OCT image dataset, making it even larger than that of techniques like light sheet microscopy. For example, imaging a entire 0.5 cm^3^ brain volume at isotropic 5 μm^3^ resolution would generate about 1 terabytes data^1^. In contrast, CAST will generate over 17 terabytes of raw data. The large volume of data, coupled with in-focus data extraction process, presents a significant challenge for the throughput of CAST image stitching. Secondly, anisotropic illumination light field results in inhomogeneous intensity distribution in stitched images. The goal of traditional stitching algorithms is to create a seamless image with uniform intensity by minimizing seam artifacts through the identification of overlap regions with minimal differences and smoothing the transitions between images^19,20^. However, when dealing with large 3D datasets stitching task, the computational cost of suppressing mosaic artifacts across all three dimensions using these methods will be substantial. Therefore, the key to improving the stitching throughput of large datasets is to find effective and computationally inexpensive methods for image intensity correction. More generally, a common challenge in stitching large datasets across imaging systems is how to fully utilize hardware resources to maximize average data throughput.

To address these challenges, we developed a fast and uniform CAST imaging system, termed *fu*CAST, together with a high-throughput image-processing pipeline for centimeter-scale samples, termed FUSION. We optimized the CAST data acquisition process and data organization format. We expanded the scanning field of view, reducing the stitching dimension from three to one to simplify the intensity correction process. We employed the hierarchical data format (HDF5) as file format and reorganized the reconstructed CAST images to many blocks to improve parallel data access speed. Moreover, we customized the peripheral component interconnect express (PCIe) storage board based on Solid-state drive (SSD) raid to achieve ultra-high input/output (I/O) bandwidth and solve the impact of I/O on program running speed. Finally, we introduced inter-frame intensity-weighted fusion to correct the intensity unevenness and achieved seamless 3D stitching. Leveraging the fuCAST imaging system together with the FUSION processing pipeline, we achieved whole-brain white-matter imaging.

## 2 Results

### 2.1 fuCAST approach for cleared brain imaging

We developed a coherent scattering imaging system that could handle intact mouse brain imaging, termed as fast and uniform clearing assisted scattering tomography (*fu*CAST). In our *fu*CAST setup (Fig.1a), a customized swept-source laser was integrated into a Michelson interferometer for coherent detection, with balanced detection implemented to suppress common-mode noise and improve sensitivity. We built sample mounting and scanning geometry in *fu*CAST system which adapted from established light-sheet microscopy configurations. An open-top sample chamber was employed to facilitate convenient sample loading and retrieval, while a scanning objective (LSM02, Thorlabs) provided lateral illumination of the specimen. For a scanning objective, the diffraction-limited two-dimensional field of view is typically constrained to be smaller than 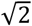 times the maximum one-dimensional scan range. To maximize utilization of the available FOV, we combined a one-dimensional galvanometric scanner with a scan lens to enable fast axial scanning, while a two-dimensional motorized translation stage provided slow-axis scanning and dynamic refocusing of the sample.

As illustrated in Fig.1a(i), the LSM02 objective provides a working distance of approximately 10.5 mm in a refractive-index-matched buffer (RI ≈ 1.46), sufficient to cover the dorsal-ventral extent of the mouse brain (∼8 mm). Although the nominal maximum scan FOV is specified as ∼5.9 mm, we experimentally observed that, under non-ideal (non-diffraction-limited) operating conditions, the objective could achieve a one-dimensional scan range of up to 12 mm, sufficient to span the medial-lateral axis of the mouse brain (∼11 mm). This extended FOV enables micron-scale resolution imaging across the entire mouse brain using commercially available scanning optics. Operating the objective beyond its diffraction-limited regime inevitably introduces field curvature and coma aberrations. Because these aberrations are well characterized, their effects were corrected computationally in the *fu*CAST pipeline using image-based aberration compensation.

By controlling photon scattering through the clearing procedure, *fu*CAST enables high resolution, cross-sectional and volumetric imaging through thick intact tissue without any contrast agent. To this end, we established a lipid-selective whole-mouse-brain clearing protocol based on extensive glutaraldehyde (GA) crosslinking, which forms a highly stable molecular network through protein-lipid and lipid-lipid crosslinks. Notably, GA can react with amine-containing lipids, such as phosphatidylethanolamine, further reinforcing lipid-associated crosslinking. This dense crosslinked network reduces tissue porosity and permeability, thereby impeding the diffusion of detergents (e.g., SDS and Triton X-100) as well as the efflux of solubilized lipid micelles. Diffusion limitations become particularly pronounced in lipid-rich regions, such as white matter, and in thick tissue samples, resulting in less efficient lipid removal in white matter compared with gray matter.

The scattering signal from cleared tissue is jointly determined by the refractive index of hydrated proteins, residual lipids, and the refractive-index-matching buffer. Based on coherent scattering theory, we simulated the scattering intensity in the *fu*CAST system as a function of local protein-to-lipid ratio under different refractive-index-matching conditions (Fig. 1b). We experimentally validated these predictions using 1-mm-thick sagittal mouse brain slices subjected to selective delipidation, and quantified signal intensities in white and gray matter across different refractive indices (Fig.1d). The mean scattering intensity of whole brain slices as a function of refractive index closely followed the simulated trend, exhibiting a minimum near RI ≈1.467. Notably, the optimal signal-to-noise contrast between white and gray matter was achieved at RI≈1.46, consistent with model predictions.

**Fig. 1.**
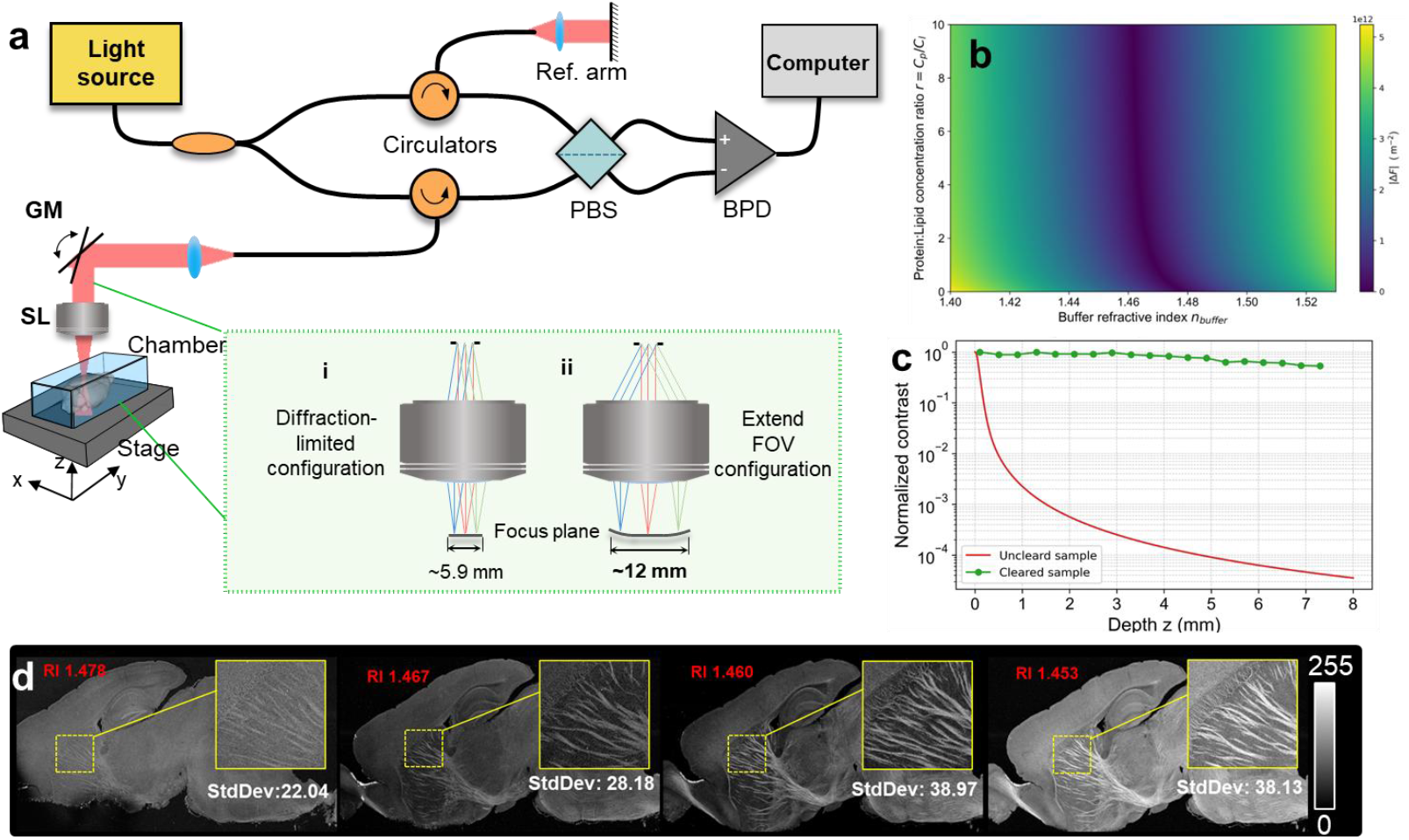
Schematic diagram of *fu*CAST system imaging for centimeter-level tissue clear sample. (a) Optical path diagram of *fu*CAST system. (b) Simulation of scattering intensity in *fu*CAST for protein-lipid complexes across varying compositions and refractive-index buffers. (c) Depth-dependent signal-to-background contrast was evaluated for cleared and uncleared samples. In uncleared tissue, signal intensity decays rapidly with increasing depth due to the dominance of multiple scattering. (d) Average signal intensity and white–gray matter contrast in 1-mm-thick sagittal brain sections after tissue clearing under different refractive-index conditions.

In addition, tissue clearing effectively mitigates the impact of multiple scattering on image formation. As shown in Fig.1c, we simulated the depth-dependent signal attenuation in uncleared tissue based on extended Huygens-Fresnel (EHF) theory^21^, under which tissue beyond a depth of approximately 0.2 mm is generally considered to be dominated by multiple scattering. In contrast, we quantified the depth-dependent signal-to-background ratio (SBR) in fully cleared whole-brain samples imaged with *fu*CAST. Across the entire brain depth, the signal exhibited less than 3 dB attenuation with increasing depth. This minimal depth-dependent decay indicates that image formation in *fu*CAST is predominantly governed by ballistic or weakly scattered photons, resulting in substantially enhanced image contrast.

### 2.2 High throughput fuCAST images processing framework

Whole-brain imaging at micron-scale resolution generates massive datasets. In *fu*CAST, these large datasets must undergo both volumetric reconstruction and multi-focal stacks stitching, making data-processing throughput a critical bottleneck for large-cohort whole-brain studies. To address this challenge, we developed a fast and uniform image-processing platform, FUSION, which leverages coordinated CPU and GPU parallelization to achieve high-throughput reconstruction and stitching of *fu*CAST datasets (Fig.2).

#### 2.2.1 Big dataset organized format

For large datasets of whole-brain images, data organization critically impacts image I/O efficiency. In FUSION, we adopt the Hierarchical Data Format version 5 (HDF5) to store fuCAST datasets, enabling efficient high-throughput data access and scalable processing. HDF5 is an open source and cross platform binary file format that supports large, complex and heterogeneous data. The specifics of unlimited on the number or size of data objects, partial I/O, and random access make HDF5 widely used in biomedical big data image storage^18^. The performance of partial I/O, i.e. reading subsets of the data, is maximized when the data selected for I/O is contiguous on disk. The blocked layout is therefore well-suited to re-slicing access to images data. To further improve I/O throughput, we employed a customized PCIe RAID controller (SSD7140A, HighPoint Technologies) configured with four solid-state drives in a disk-stripping array (RAID-0). This configuration saturates the ×16 PCIe 3.0 bus bandwidth, providing sustained data transfer rates exceeding 14 GB/s while supporting up to 64 TB of storage capacity. Compared with conventional mechanical hard drives, this setup delivers more than two orders of magnitude improvement in I/O performance.

For *fu*CAST data, unlike other scanning imaging methods which acquire the sample information point by point, *fu*CAST imaging the sample is alone the A-scan, results in the raw data are A-line by A-line storage. In the *fu*CAST reconstruction data reorganization, we store the whole-brain data in a group in HDF5, each stack as a dataset individually, and each dataset is divided into blocks based on the computer’s memory size and CPU cores number (Fig. 2c). Sequential storage of each block and CPU multi-threaded parallel reads can fully increase the throughput of data reading and processing.

**Fig. 2.**
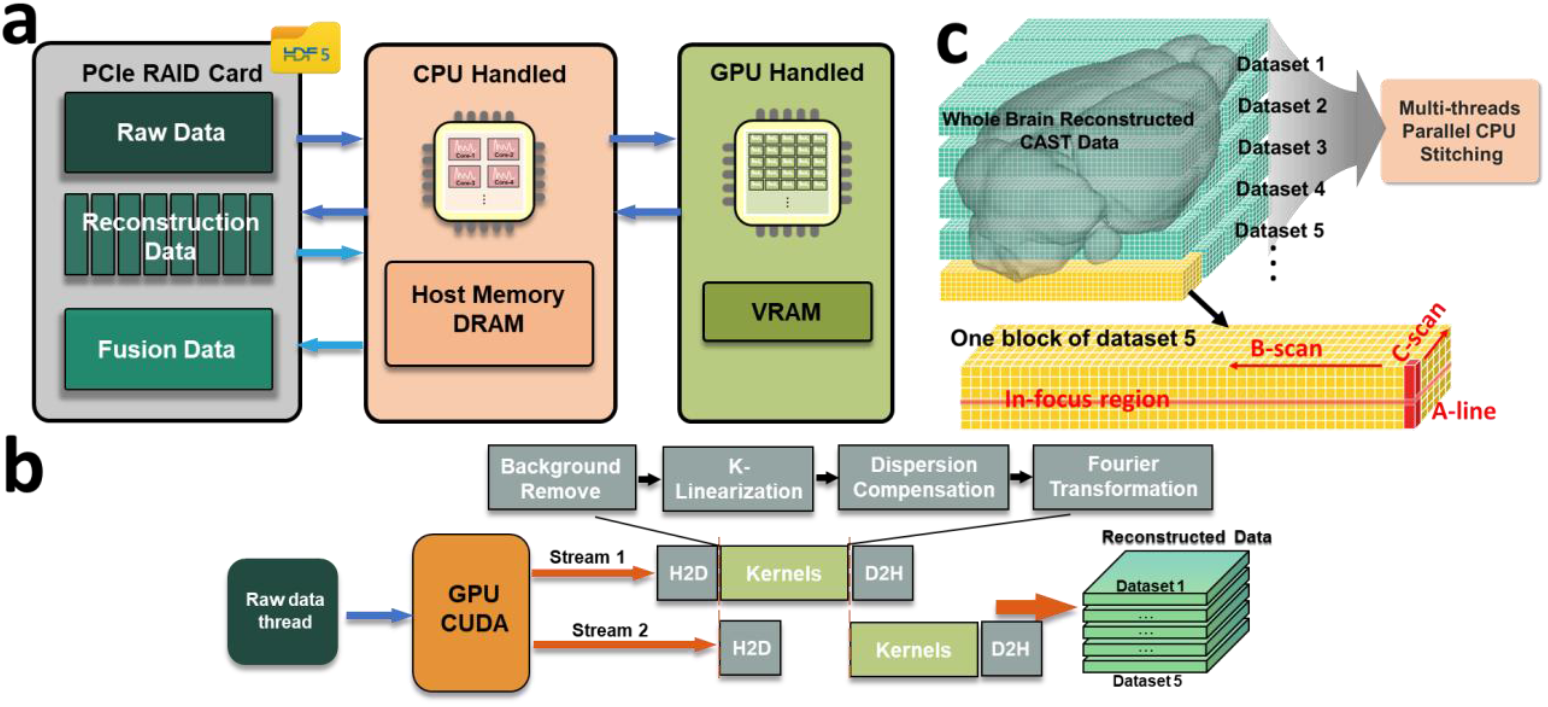
Big dataset processing and reorganization pipeline for *fu*CAST images. (a) A dataflow architecture for GPU-accelerated reconstruction and CPU-parallel stitching enabled by a custom PCIe SSD array. (b) HDF5-based blocked storage of reconstructed datasets. (c) Schematic of multithreaded GPU-accelerated pipeline.

#### 2.2.2 High-throughput data flow of image reconstruction and stitching pipeline

Reconstruction and stitching impose fundamentally different demands on hardware performance in *fu*CAST data processing. Image reconstruction involves operations such as dispersion compensation and repeated Fourier transforms, making it a compute-intensive task that benefits primarily from GPU acceleration. In contrast, large-scale volumetric stitching is dominated by data movement rather than computation. Because the imaging field of view is extended in *fu*CAST, three-dimensional stitching can be effectively reduced to one-dimensional concatenation, eliminating computationally expensive overlap alignment and registration steps. As a result, stitching becomes an I/O-intensive task whose performance is primarily limited by data throughput.

Motivated by this distinction, we designed a heterogeneous data-processing framework that combines GPU-parallel acceleration for reconstruction with high-throughput, CPU-based stitching centered on a PCIe RAID storage architecture (Fig.2a). GPU-parallel processing in FUSION is implemented using a producer-consumer paradigm, with dedicated threads for data loading, transfer, and computation to form a fully pipelined execution workflow (Fig.2b). As illustrated in Fig. 2c, reconstructed data are partitioned into spatial chunks and stored as separate datasets. During stitching, each CPU thread sequentially reads chunked datasets from different stacks along the A-line direction, extracts and fuses the in-focus regions following the procedure described in Section 2.3 and generates a complete A-block. The resulting block is then written on the FUSION dataset, after which the thread proceeds to the next spatial position. Block sizes are optimized during reconstruction to maximize utilization of available CPU cores and system memory, thereby improving overall stitching throughput.

### 2.3 High throughput whole-brain image stitching and quantitative evaluation

#### 2.3.1 Focus plan and strip correction

As shown in Fig. 3a, extending the field of view introduces field curvature. In FUSION, this effect is corrected by extracting the in-focus regions across the volume. Specifically, a subset of in-focus positions within the sample is identified and used to fit a three-dimensional curved focal surface. This fitted surface captures both the intrinsic field curvature of the extended FOV and small optical path length shifts introduced during slow-axis scanning. During in-focus extraction, the A-line length is kept constant across lateral coordinates, despite focal-plane bending shifting the focal coordinates in depth. For efficient matrix-based computation, the extracted three-dimensional in-focus volume is flattened into a planar representation. After stack-wise intensity correction, the in-focus A-line data are remapped back to their original Z-stitching coordinates, restoring the curved focal surface.

**Fig. 3.**
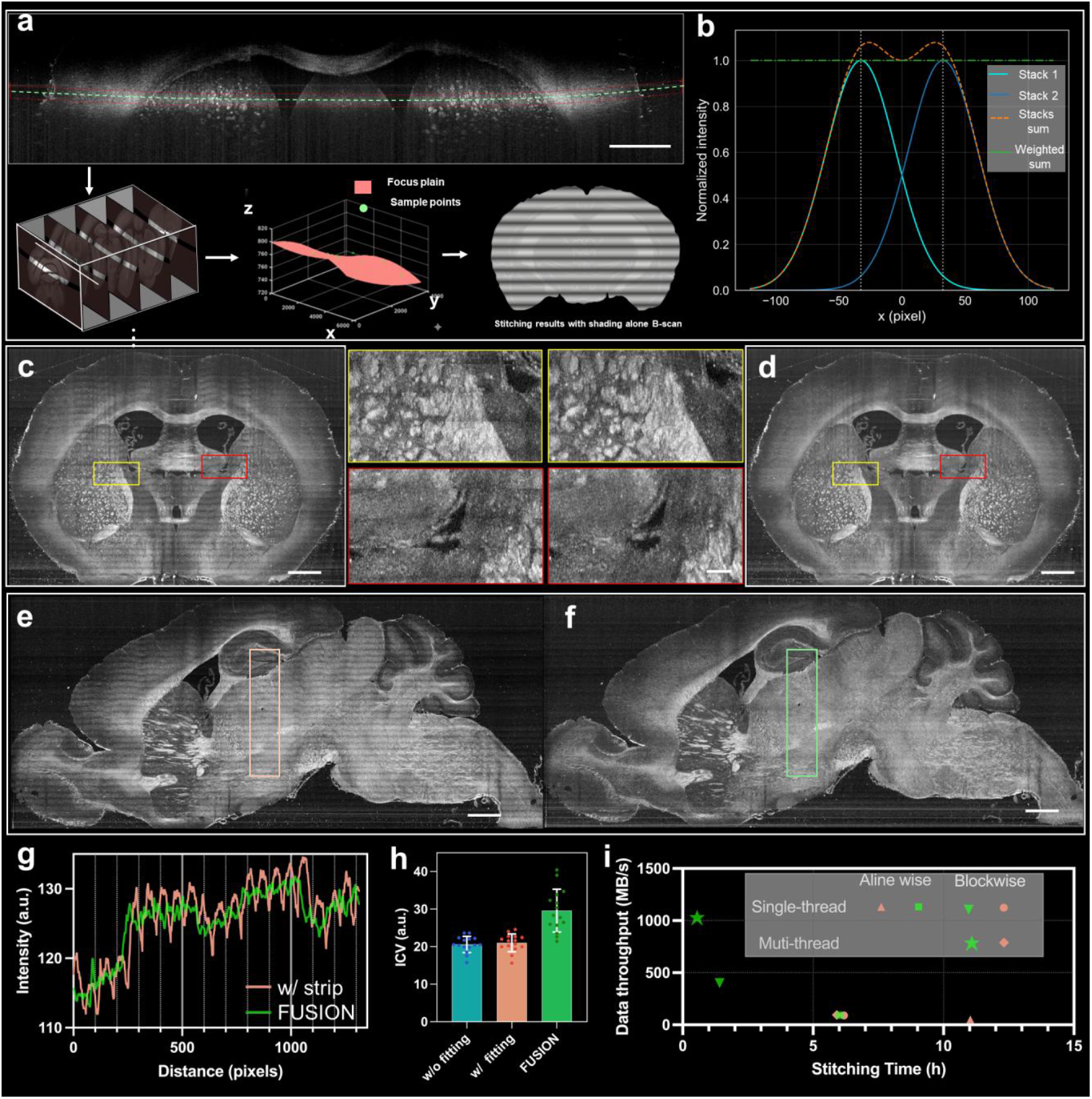
Whole-brain images stitching and quantitative performance evaluation using FUSION. (a) Focal-plane fitting and in-focus image extraction of field-curvature correction. (b)Dual-Gaussian weighted fusion of adjacent stacks for striping artifact correction. (c-d)Comparison between stitching results obtained without focal-plane fitting and FUSION results. (e-f) Comparison of stitching results without intensity striping correction and with the FUSION method. (g) The intensity profiles along the area within the rectangles in the stripes demonstrate that FUSION corrects the intensity fluctuations more effectively, while preserving tissue characteristic information. (h) The inverse coefficient variation (ICV) further quantitatively evaluates the destripe ability. A larger ICV corresponds to the flatter intensity profile and indicates a better correction quality. (i) Impact of different stitching strategies and hardware configurations on data throughput. The FUSION approach, indicated by green star, achieves the highest throughput.

Due to a Gaussian beam exhibiting axial intensity variation and coherent scattering imaging is linear, direct stitching of in-focus images introduces striping artifacts caused by intensity discontinuities between adjacent stacks. These artifacts degrade image quality and adversely affect downstream analyses such as white-matter tract tracing and quantitative measurements. To mitigate this effect, FUSION incorporates an inter-stack fusion strategy (Fig. 3b). During in-focus extraction, an axial range corresponding to twice the full width at half maximum (FWHM) of the Gaussian profile is selected, resulting in a 50% overlap between adjacent stacks. The overlapping regions are then combined using weighted averaging, yielding a smooth and intensity-uniform image that effectively suppresses striping artifacts.

#### 2.3.2 Performance evaluation of stitching algorithm

Extraction of in-focus regions based on the fitted focus plain and image intensity correction operations improve image uniform and effectively reduce the stitching blinding effect. Fig. 3d and Fig. 3f show coronal and sagittal cross sections of the whole brain 3D image, respectively. The structure of the mice brain is intact, the in-focus regions are extracted accurately, and the blinds effect is significantly suppressed after intensity correction. This result demonstrates the effectiveness of our stitching method. We compared the FUSION stitching results with results which performed without field-curvature correction or without intensity striping correction (Fig.3c,e).

A curved in-focus band along the B-scan direction necessitates focal-surface fitting for consistent in-focus extraction. In addition, small optical path length variations introduced during C-scan translation can shift equal-optical-path points, leading to noticeable inter-stack misalignment (Fig.3c). And inaccuracies in focal-plane estimation render intensity correction ineffective and can even exacerbate striping artifacts. The enlarged images of the yellow and red box regions stitched by FUSION method shown that the fine nerve fiber bundles are structurally intact, and the blind effect and misalignment of the images brought about by stitching are virtually nonexistent over a range crossing more than five stacks. In addition, if the stacks are not intensity corrected, the stitching result will produce obvious strip artifacts (Fig. 3e). In contrast, after applying the inter-stack intensity-weighted fusion in FUSION, the reconstructed images exhibit highly uniform intensity, with no observable striping artifacts across the entire sagittal section view (Fig. 3f).

To quantitatively assess correction performance, we analyzed intensity profiles extracted from a defined region of interest (ROI), corresponding to the boxed area in the sagittal section (Fig. 3g). In the uncorrected images, these profiles exhibit pronounced intensity variations when traversing from non-striped regions into striped regions. In contrast, FUSION effectively suppresses such intensity fluctuations while preserving intrinsic tissue features, outperforming the other correction strategies. For further quantitative validation, we employed the inverse coefficient of variation (ICV) as a metric to evaluate de-striping performance^22^. Among all methods tested, FUSION achieved the highest ICV values, indicating superior stripe suppression (Fig. 3h).

We further evaluated the throughput of FUSION in comparison with other stitching methods. Fig. 3i shows the comparison of program execution time and overall program throughput among different stitching measurements. All tests were based on the same whole mice brain dataset. We measured the execution time and data throughput of the A-line-based and our proposed block-based approach on HDD and SSD array RAID card, respectively. Finally, the stitched A-line is saved and the next Aline stitching task is started. When using HDD, Aline-wise method would take more than 11 hours to complete image stitching of a whole mouse brain. This would far exceed the image acquisition time, severely reducing the data throughput of the entire process of acquisition, reconstruction and processing in the *fu*CAST system. Only 50.96 MB/s is not entirely caused by the slow I/O speed of the HDD. The Aline-wise stitching algorithm less than doubled the throughput even when replaced with an SSD array card with over 13GB/s I/O speed. In other words, Aline-wise solution will not maximum utilization of I/O throughput and CPU computational efficiency. The blocked stitching strategy adopted in FUSION will not largely increase the stitching speed when using HDD. The speed is increased by less than factor one. Moreover, the speeds of single-threaded and multi-threaded programs on the CPU are basically the same, which indicates that the upper limit of the I/O speed of the HDD limits the performance of the program. After replacing the HDD with a higher bandwidth SSD array RAID, the CPU single-threaded stitching algorithm speed will be about 4.3 times faster than the Aline-wise algorithm, and the speed will be about 10.95 times faster with CPU multi-threading. It will take only 0.55 hours to complete the stitching of the entire mouse brain dataset. Based on our blocked stitching algorithm, the arithmetic power of CPU and the bandwidth advantage of SSD can be fully utilized. Stitching throughput exceeds 1 GB/s in our hardware configuration. For different computers, we can optimize the size of the blocks and the number of parallel CPU threads to achieve the highest stitching throughput of the computer. It is worth mentioning that the throughput will not decrease as the dataset grows larger, e.g., for larger macaque brain or human brain datasets.

#### 2.3.3 fuCAST enables visualization of entirely myelinated fiber tracts

Our imaging method allows the B-scan field of view to reach more than 12 mm and cover the cranial width of mice brain. While the telecentric sacrifice increases the complexity of our stitching algorithm, the large field of view also reduces the dimensionality and number of stitches. One-dimensional stitching along the Z-direction provides us with unstitched enface section images (Fig. 4a). The direction of nerve fibers throughout the brain and the interconnections between different brain regions can be shown in one field of view with the same signal-to-noise rate. And the distribution of nerve fibers is clearly shown in the magnified image of the corresponding cortical region in Fig. 4b, which nerve fibers exhibit stronger scattering than other tissues after tissue clear. The connections of nerve fibers on a whole-brain scale can be more fully demonstrated in the 3D rendered image of Fig. 4c. This demonstrates the great potential of the *fu*CAST method for applications in full-brain connectivity mapping as well as neuroscience research on diseases such as stroke.

**Fig. 4.**
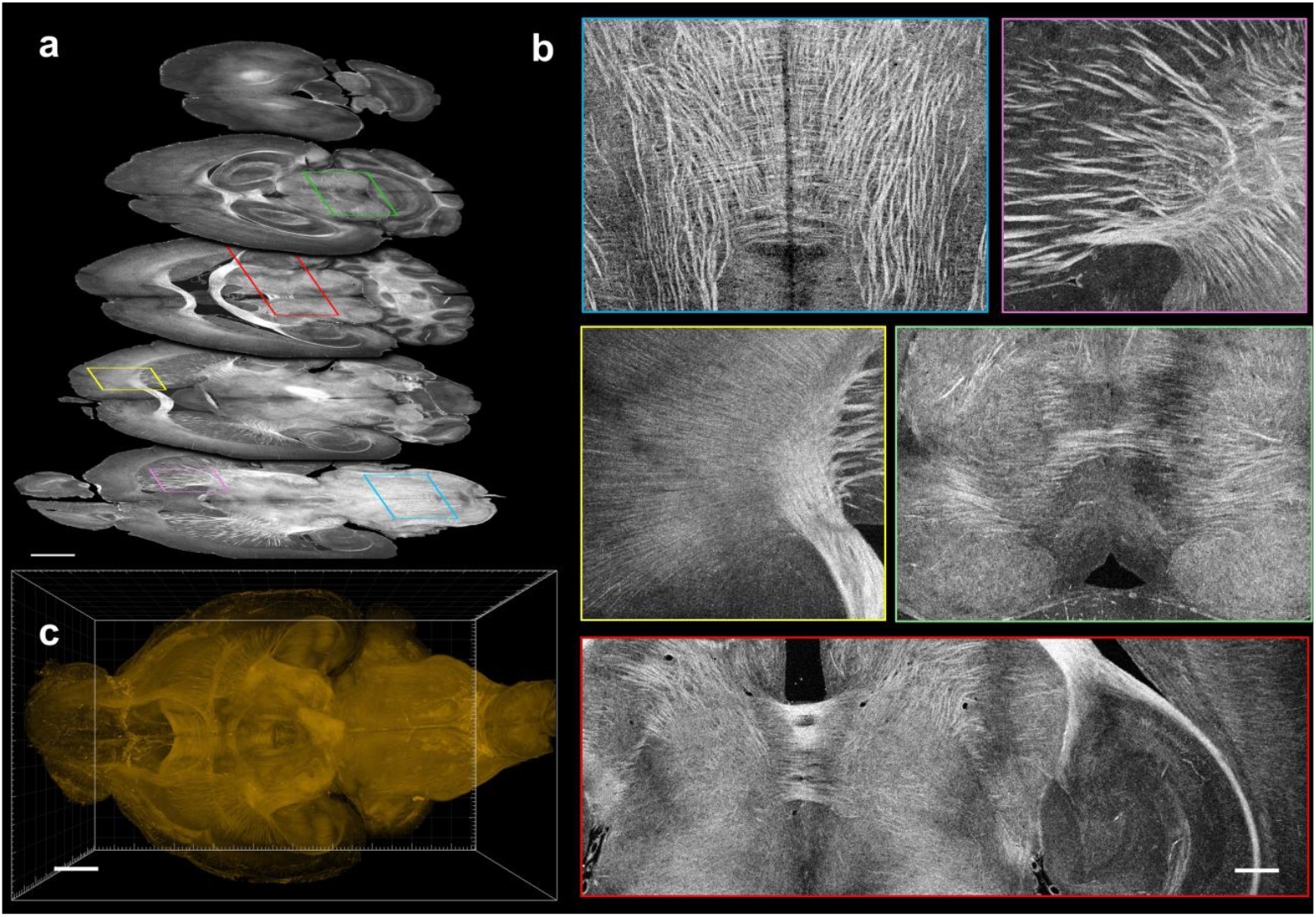
*fu*CAST enables visualization of entirely myelinated fiber tracts (a) En face images acquired at different depths demonstrate robust white-matter contrast across multiple brain regions. Axial-plane images are acquired as continuous views without stitching due to the extended field of view in fuCAST. (b) Magnified views of regions of interest from different brain regions in (a). (c) 3D visualization of white-matter distribution across the whole brain.

## 3 Discussion

Coherent scattering imaging holds broad promise for biomedical imaging due to its ultra-high information throughput and sensitivity. However, its application to whole-brain imaging has remained limited, largely owing to insufficient optical penetration in intact tissue. Here, by integrating tissue clearing with coherent scattering imaging, we develop fuCAST, an intact-brain coherent scattering imaging approach, together with a high-throughput data-processing platform, FUSION. This framework enables selective lipid removal across the whole brain to enhance white-matter scattering contrast, allowing comprehensive mapping of whole-brain white-matter organization.

The *fu*CAST approach increases system throughput and reduces stitching dimensionality by extending the effective field of view of a commercial objective. In post-processing, focal-surface fitting and in-focus extraction are employed to compensate for field curvature, yielding robust stitching performance. We acknowledge, however, that operating the objective beyond its diffraction-limited regime inevitably introduces off-axis aberrations, such as coma, in addition to field curvature. In the present work, these higher-order aberrations were not further corrected, as no obvious image degradation was observed toward the edges of the extended field of view. Because such aberrations are well characterized, image-based deconvolution could, in principle, further mitigate their impact on image quality, albeit at the cost of substantially increased computational burden. Alternatively, integrating adaptive optics to correct these aberrations during image acquisition represents a promising direction for future system optimization.

We developed the FUSION framework to address throughput limitations in image reconstruction and stitching. We note that the raw reconstruction speed achieved in this work is not the theoretical maximum, and further acceleration could be obtained by adopting higher-performance GPUs. However, the primary contribution of FUSION lies not in maximizing a single processing stage, but in integrating the entire image-processing workflow into a unified, high-throughput framework. At the core of this framework is a customized high-bandwidth SSD RAID I/O architecture, which enables efficient coordination between reconstruction, stitching, and data movement. By alleviating I/O bottlenecks and balancing compute- and I/O-intensive tasks, FUSION achieves robust end-to-end throughput for large-scale CAST datasets. In our system, the SSD RAID card achieves a speed of 14 GB/s. However, the latest HighPoint RocketAIC 7608A series NVMe AIC SSDs harness eight top-tier PCIe Gen5 or Gen4 M.2 SSDs combined with HighPoint’s revolutionary PCIe Switch Architecture and industry-leading RAID technology. This enables real-world transfer speeds of up to 56 GB/s and storage capacities reaching 64 TB, all from a single PCIe Gen5 x16 slot. Our algorithm could dynamically adjust the data block size based on hardware specifications to maximize stitching throughput for the given configuration. More importantly, the throughput of this block-based algorithm remains unaffected by the size of the sample dataset. While the hardware configurations employed in this paper only enable stitching the centimeter-scale samples, such as mouse brain, we are confident that using higher-performance SSD RAID cards will enable faster speeds and the ability to handle larger sample stitching tasks within the framework of our parallel stitching algorithm. Additionally, servers equipped with more CPU cores and larger memory capacities offer significant potential for further speed improvements. In future imaging of tissue-cleared human brain samples, the CAST technology combined with our stitching algorithm will demonstrate significant throughput advantages.

Finally, the methods presented in this work extend coherent scattering imaging beyond its traditional use in clinical ophthalmologic and endoscopic studies to broaden biomedical imaging applications. The tissue clearance-assisted scattering imaging methods have enabled label free imaging of large *ex vivo* samples such as mouse tissue. For *fu*CAST images of mouse brain, the connections of nerve fibers throughout the brain can be clearly observed, which will provide a more convenient tool for the study of Alzheimer’s disease, stroke, and brain connectivity mapping. Our proposed high throughput splicing collocation method for large datasets will further improve the imaging throughput of the CAST method and facilitate its application in scenarios such as drug screening.

## 4 Material and Methods

### Tissue preparation and mounting

Tissue preparation was performed using a SWITCH-based clearing protocol, selected for its controllable lipid removal and preservation of native biomolecules, which enables subsequent immunostaining. Brains from young adult wild-type C57BL/6 mice (6-8 weeks) were harvested following transcranial perfusion with 20 mL ice-cold PBS, followed by 20 mL of ice-cold fixative containing 4% paraformaldehyde (PFA) and 1% glutaraldehyde (GA) in 1× PBS. Excised brains were post-fixed in the same perfusion solution for 3 days at 4 °C with gentle agitation. Samples were then washed twice in PBST for 6 h each at room temperature, followed by overnight incubation in an inactivation solution (1× PBS, 4% (w/v) acetamide, and 4% (w/v) glycine) at 37 °C. After two additional washes (6 h each) in clearing solution containing 200 mM sodium dodecyl sulfate (SDS) and 20 mM sodium sulfite, tissues were incubated in the same clearing solution in a 70 °C water bath for lipid removal. Clearing duration and temperature were adjusted according to sample size to achieve different levels of optical transparency.

Because water-based cleared tissues are mechanically soft, samples were embedded in acrylamide hydrogel to provide mechanical support during imaging. Briefly, 1.2 mL of 40% acrylamide solution (Bio-Rad) was diluted with 4.8 mL of exPROTOS solution and thoroughly mixed. Polymerization was initiated by adding 30μL ammonium persulfate (APS) and 6μL N,N,N’,N’-tetramethyl ethylenediamine (TEMED). The mixture was immediately poured into a 3D-printed mold containing the cleared brain tissue, ensuring complete encapsulation. After gelation at room temperature, the embedded samples were transferred to the final refractive-index–matching buffer and incubated on a shaker at room temperature for 24 h to allow full equilibration. The samples were then mounted in a custom holder for fuCAST imaging.

### Intact mouse brain fuCAST images acquisition

In fuCAST system includes three scanning directions, which are commonly known as A-Scan, B-Scan and C-Scan. As shown in Fig.1a, a two axes long period motorized translation stage combined galvo mirror (Cambridge Technology Inc., D05822) achieves a big FOV’s 3D scanning imaging. The scan lens (Thorlabs Inc., LSM02) employed as objective lens and resulted in a working distance of about 6.7 mm and a later resolution of ∼8 μm. This scanning objective has only a small field of view, but it is difficult to find commercialized objectives that can satisfy the large field of view and high-resolution scanning objective for whole-brain imaging in mice. Therefore, we still used the commonly used Telecentric Scanning Lenses (LSM02, Thorlabs, FOV 4.7 x 4.7 mm^2^) in the fuCAST method, which was experimentally found to achieve a single directional scanning range of about 12.5 mm. In addition, instead of using a galvanometer for the slow axis, we chose a long period translation stage, which ensures that the maximum XY field of view is sufficient to cover the entire axial section of the mouse. A homemade wavelength swept laser was employed to realize A-Scan utilized a semi-conductor optical amplifier (Covega Corp., BOA-4379) as the gain medium and a polygon scanner (Lincoln Laser Co., SA34/DT-72-250-025-AA/01B) as the tunable filter to rapidly tune the wavelength at a rate of 54 kHz. This source had a center wavelength of 1300 nm and a sweeping range of 110 nm, resulting in an axial resolution of 9 μm in air. The axial Rayleigh length of a Gaussian beam at this girdle diameter is only about 150 μm.

### Hardware features

To realize the storage and high-speed processing of huge amounts of CAST 3D brain imaging data, our reconstruction and stitching algorithms were built on a high-performance data processing workstation with a 128 GB physical Memory (RAM). All the subsequent experimental results in this article are based on this workstation. For CAST system, an 8 mm × 12 mm × 20 mm volume of tissue clear mouse brain will produce approximately 6.7 TB raw data. As shown in Fig. 1c, the entire mouse brain will have about 54 times more data than the physical memory of the workstation. In the face of this difference between the two orders of magnitude of data volume and processing power, it is necessary to increase the throughput of data processing by increasing the speed of reading and writing data. In addition, a single SSD cannot handle such a huge amount of data. Therefore, in our system, we customized a PCIe Gen3 card (High Point Technologies Inc., SSD7140A) with 4 solid-state drives (SSDs) array based on disk striping (RAID 0). The board allows customers to saturate x16 lanes of PCIe 3.0 bus-bandwidth with sustained transfer performance over 14 GB/s while supporting up to 64 TB of storage. Moreover, we employ an 18-Cores CPU (Intel corp., Core™X i9-9980XE) and a GPU (NVIDIA Corp., RTX™ 2070, 8G) for parallel computation to further accelerate the throughput of stitching.

## Acknowledgements

This research was funded by National Institutes of Health (NIH) (P41EB-015903) and (R00AG059946)

## References

1. Glaser, A.K., et al., 2022. A hybrid open-top light-sheet microscope for versatile multi-scale imaging of cleared tissues. Nat Methods 19, 613–619. 10.1038/s41592-022-01468-5

2. Yamashita, K. et al. A whole-brain single-cell atlas of circadian neural activity in mice. Science 0, eaea3381 (2025).

3. Yun, D. H. et al. Uniform volumetric single-cell processing for organ-scale molecular phenotyping. Nat Biotechnol 1–12 (2025) doi:10.1038/s41587-024-02533-4.

4. Jing, D. et al. Tissue clearing of both hard and soft tissue organs with the PEGASOS method. Cell Res 28, 803– 818 (2018).

5. Schneidereit, D. et al. An advanced optical clearing protocol allows label-free detection of tissue necrosis via multiphoton microscopy in injured whole muscle. Theranostics 11, 2876–2891 (2021).

6. Wei, M. et al. Volumetric chemical imaging by clearing-enhanced stimulated raman scattering microscopy. Proceedings of the National Academy of Sciences 116, 6608–6617 (2019).

7. Draelos, M. et al. Contactless optical coherence tomography of the eyes of freestanding individuals with a robotic scanner. Nat Biomed Eng 5, 726–736 (2021).

8. Ughi, G. J. et al. A neurovascular high-frequency optical coherence tomography system enables in situ cerebrovascular volumetric microscopy. Nat Commun 11, 3851 (2020).

9. Wang, H., Zhu, J. & Akkin, T. Serial optical coherence scanner for large-scale brain imaging at microscopic resolution. NeuroImage 84, 1007–1017 (2014).

10. Lefebvre, J., Castonguay, A., Pouliot, P., Descoteaux, M. & Lesage, F. Whole mouse brain imaging using optical coherence tomography: reconstruction, normalization, segmentation, and comparison with diffusion MRI. NPh 4, 041501 (2017).

11. Yuan, W. et al. Theranostic OCT microneedle for fast ultrahigh-resolution deep-brain imaging and efficient laser ablation in vivo. Science Advances 6, eaaz9664 (2020).

12. Ren, J., Choi, H., Chung, K. & Bouma, B. E. Label-free volumetric optical imaging of intact murine brains. Sci Rep 7, 46306 (2017).

13. Jian, Y., Wong, K. & Sarunic, M. V. Graphics processing unit accelerated optical coherence tomography processing at megahertz axial scan rate and high resolution video rate volumetric rendering. JBO 18, 026002 (2013).

14. Zabic, M., Matthias, B., Heisterkamp, A. & Ripken, T. Open source optical coherence tomography software. Journal of Open Source Software 5, 2580 (2020).

15. Wang, M. et al. Physics-guided deep learning-based real-time image reconstruction of fourier-domain optical coherence tomography. Biomed. Opt. Express, BOE 15, 6619–6637 (2024).

16. Bria, A. & Iannello, G. TeraStitcher - A tool for fast automatic 3D-stitching of teravoxel-sized microscopy images. BMC Bioinformatics 13, 316 (2012).

17. Bria, A., Bernaschi, M., Guarrasi, M. & Iannello, G. Exploiting Multi-Level Parallelism for Stitching Very Large Microscopy Images. Front Neuroinform 13, 41 (2019).

18. Hörl, D. et al. BigStitcher: Reconstructing high-resolution image datasets of cleared and expanded samples. Nat Methods 16, 870–874 (2019).

19. Peng, T. et al. A BaSiC tool for background and shading correction of optical microscopy images. Nat Commun 8, 14836 (2017).

20. Wang, S. et al. A deep learning-based stripe self-correction method for stitched microscopic images. Nat Commun 14, 5393 (2023).

21. Untracht, G. R. et al. Spatially offset optical coherence tomography: Leveraging multiple scattering for high-contrast imaging at depth in turbid media. Sci. Adv. 9, eadh5435 (2023).

22. Pollatou, A. An automated method for removal of striping artifacts in fluorescent whole-slide microscopy. Journal of Neuroscience Methods 341, 108781 (2020).

